# Effects of enriched housing on the neuronal morphology of mice that lack zinc transporter 3 (ZnT3) and vesicular zinc

**DOI:** 10.1101/754747

**Authors:** Brendan B. McAllister, Sarah E. Thackray, Brenda Karina Garciá de la Orta, Elise Gosse, Purnoor Tak, Colten Chipak, Sukhjinder Rehal, Abril Valverde Rascón, Richard H. Dyck

## Abstract

In the central nervous system, certain neurons store zinc within the synaptic vesicles of their axon terminals. This vesicular zinc can then be released in an activity-dependent fashion as an intercellular signal. The functions of vesicular zinc are not entirely understood, but evidence suggests that it is important for some forms of experience-dependent plasticity in the brain. The ability of neurons to store and release vesicular zinc is dependent on expression of the vesicular zinc transporter, ZnT3. Here, we examined the neuronal morphology of mice that lack ZnT3. Brains were collected from mice housed under standard laboratory conditions and from mice housed in enriched environments – large, multilevel enclosures with running wheels, numerous objects and tunnels, and a greater number of cage mates. Golgi-Cox staining was used to visualize neurons for analysis of dendritic length and dendritic spine density. Neurons were analyzed from the barrel cortex, striatum, basolateral amygdala, and hippocampus (CA1). ZnT3 knockout mice, relative to wild type mice, exhibited increased basal dendritic length in the layer 2/3 pyramidal neurons of barrel cortex, independently of housing condition. Environmental enrichment decreased apical dendritic length in these same neurons and increased dendritic spine density on striatal medium spiny neurons. Elimination of ZnT3 did not modulate any of the effects of enrichment. Our results provide no evidence that vesicular zinc is required for the experience-dependent changes that occur in response to environmental enrichment. They are consistent, however, with recent reports suggesting increased cortical volume in ZnT3 knockout mice.

## 1.1 INTRODUCTION

Zinc is an essential component of all cells (Chasapis et al., 2012), including those that compose the central nervous system (CNS)^*^. Much of this zinc is tightly bound in protein structures (Vallee & Auld, 1990; Andreini et al., 2006), but within certain regions of the brain, a considerable portion (∼20%) is found in a “free” (i.e., unbound or loosely-bound) state within the synaptic vesicles of neurons (Pérez-Clausell & Danscher, 1985; Cole et al., 1999). This vesicular zinc can be released into the synaptic cleft in an activity-dependent manner (Assaf & Chung, 1984; Howell et al., 1984; Aniksztejn et al., 1987; Li et al., 2001; Ueno et al., 2002), where it can act as a neuromodulator at many membrane-bound receptors or enter cells to affect second messenger systems (reviewed by McAllister & Dyck, 2017). Histochemical methods for visualizing free zinc reveal that zinc-containing axon terminals are abundant in the neocortex, hippocampus, amygdala, and striatum, among other regions (Danscher et al., 1982; Pérez-Clausell & Danscher, 1985; Frederickson et al., 1992). In the neocortex and hippocampus, vesicular zinc is found specifically within the terminals of glutamatergic neurons (Beaulieu et al., 1992; Sindreu et al., 2003), indicating that zinc and glutamate are co-released.

The function of vesicular zinc in the brain has been studied using mice in which zinc transporter 3 (ZnT3), the transporter responsible for storing zinc in synaptic vesicles (Palmiter et al., 1996; Wenzel et al., 1997), has been genetically deleted. Accordingly, these ZnT3 knockout (KO) mice show a complete lack of vesicular zinc (Cole et al., 1999; Linkous et al., 2008). While this does not appear to cause gross behavioural abnormalities (Cole et al., 2001; but see Yoo et al., 2016), it does affect several functions, including fear memory (Martel et al., 2010, 2011), object recognition memory (Martel et al., 2011), contextual discrimination (Sindreu et al., 2011), skilled motor learning (Thackray et al., 2017), and both texture and auditory frequency discrimination (Wu & Dyck, 2018; Kumar et al., 2019). In addition, there is some evidence that ZnT3 KO mice exhibit abnormalities in brain morphology. A recent study indicated that the size of the neocortex is enhanced in ZnT3 KO mice, as is the neocortical expression of markers for neurons and neuronal processes (Yoo et al., 2016). Consistent with this, analysis by *ex vivo* magnetic resonance imaging (MRI) showed that the volume of the parietal cortex and corpus callosum is increased in ZnT3 KO mice (McAllister et al., 2018). However, to our knowledge, morphological abnormalities at the level of individual neurons have yet to be examined.

Vesicular zinc may also be involved in some forms of experience-dependent plasticity. We and others have shown that vesicular zinc levels in the barrel cortex are regulated by sensory experience, with levels increasing in response to sensory deprivation (e.g., by whisker plucking) and decreasing in response to repeated whisker stimulation (Czupryn & Skangiel-Kramska, 2001; Brown & Dyck, 2002, 2005; Nakashima & Dyck, 2010). While it has not been conclusively determined whether these changes are a causal factor in the cortical plasticity that can also occur in response to sensory experience (Glazewski & Fox, 1996; Bender et al., 2006), the correlation between the two phenomena indicates a possible role for zinc. Furthermore, evidence from experiments on the auditory system also supports the conclusion that vesicular zinc is regulated by sensory experience and that this regulation has consequences for synaptic function. In the dorsal cochlear nucleus, sustained loud sound exposure reduces vesicular zinc levels and evoked zinc release from the parallel fibers, and these changes are associated with a loss of zinc-induced inhibition at parallel fiber synapses (Kalappa et al., 2015). Finally, other results have shown vesicular zinc to be involved in a separate form of experience-dependent plasticity: adult hippocampal neurogenesis. Specifically, ZnT3 KO mice do not exhibit an increase in neurogenesis in response to hypoglycemia, whereas wild type (WT) mice do (Suh et al., 2009).

In the present experiment, we used Golgi-Cox staining to visualize neurons, and we conducted a morphological analysis of neurons in several brain regions that are rich with vesicular zinc, comparing WT and ZnT3 KO mice, and also assessing how neuronal morphology is altered when mice were exposed to environmental enrichment. We sought to address two main questions. 1) Does a lack of vesicular zinc cause abnormalities in neuronal morphology? This question was of particular interest given recent evidence that neocortical volume is increased in ZnT3 KO mice (Yoo et al., 2016; McAllister et al., 2018). 2) Do ZnT3 KO mice exhibit experience-dependent changes in neuronal morphology in response to environmental enrichment? Such effects of enrichment are well-documented, primarily in rats (Greenough & Volkmar, 1973; Greenough et al., 1973; Uylings et al., 1978; Kolb et al., 2003a, 2003b; Gelfo et al., 2009), but also in mice (Rampon et al., 2000; Faherty et al., 2003; Restivo et al., 2005; Karelina et al., 2012; Jung & Herms, 2014). Given the deficiencies in experience-dependent plasticity described above, we hypothesized that ZnT3 KO mice, relative to WT mice, would show fewer or smaller effects of environmental enrichment on neuronal morphology, both in terms of dendritic branching and spine density.

## 1.2 MATERIAL AND METHODS

### 1.2.1 Animals

All protocols were approved by the Life and Environmental Sciences Animal Care Committee at the University of Calgary and followed the guidelines for the ethical use of animals provided by the Canadian Council on Animal Care. The mice used in these experiments were housed in temperature- and humidity-controlled rooms maintained on a 12:12 light/dark cycle (lights on during the day). Mice were provided with food and water *ad libitum*. WT and ZnT3 KO mice, on a mixed C57BL/6×129Sv background, were bred from heterozygous pairs. Offspring were housed in standard cages (described in the next section) with both parents until they were weaned at 3-weeks of age. Only female offspring were used in this experiment.

### 1.2.2 Environmental Enrichment

At 4-weeks of age, mice were either transferred from standard housing to enriched housing (WT: *n* = 8; ZnT3 KO: *n* = 7) or remained in standard housing (WT: *n* = 6; ZnT3 KO: *n* = 6). Standard housing consisted of a cage (28 × 17.5 × 12 cm) with bedding, nesting material, one enrichment object, and 2-5 same-sex littermates. Enriched housing consisted of a much larger cage (60 × 60 × 60 cm) with two stories that the mice could climb between. In addition to bedding and nesting material, these cages contained tunnels, running wheels, multiple enrichment objects, and multiple sources of food. To increase novelty, the objects and food sources were re-positioned weekly, and some objects were replaced with different ones. To provide social enrichment, mice were housed in larger groups of 8-10, including littermates as well as similarly aged non-littermates. At 12-weeks of age the mice were killed, and brains were collected for Golgi-Cox staining.

### 1.2.3 Golgi-Cox Staining

The Golgi-Cox method allows for the impregnation of neurons throughout an intact rodent brain, and subsequent visualization of these cells, including dendritic processes and spines, in histological sections. It provides an inexpensive means of visualizing neurons with good resolution across multiple regions throughout the brain, which was ideal for the present experiment. Importantly, only a subset of neurons in a given region are stained by this method, making it highly useful for conducting morphological analysis of whole neurons that are relatively unobscured by other cells. Golgi-Cox staining was conducted following the protocol of Gibb and Kolb (1998), with minor modifications for staining the brains of mice. Mice were deeply anesthetized with sodium pentobarbital, then perfused transcardially with 100 ml of 0.9% saline. Brains were extracted, weighed, wrapped in gauze, and placed in jars containing 20 ml of Golgi-Cox solution (Glaser & van der Loos, 1981), where they were stored for 10 days in the dark. Brains were then transferred to a 30% sucrose solution and stored for 3-7 days in the dark, before being cut into 250 •m coronal sections using a vibratome (Leica VT1000S). The sections were pressed onto 1.5% gelatin-coated slides and stored in a humidity chamber for 2-5 days prior to staining. The slides were processed for staining as follows: 1 min in distilled water (dH_2_0); 30 min in ammonium hydroxide (in the dark); 1 min in dH_2_0; 30 min in Kodak Fix for Film diluted 1:1 with dH_2_0 (in the dark); 1 min in dH_2_0; 1 min in 50% ethanol; 1 min in 70% ethanol; 1 min in 95% ethanol; 3 × 5 min in 100% ethanol; 10 min in a solution of 1/3 xylenes, 1/3 chloroform, 1/3 ethanol; 2 × 15 min in xylenes. Finally, slides were cover-slipped with Permount.

### 1.2.4 Morphological Analysis

For analysis of dendritic morphology, neurons were selected and traced from each of the following regions of interest: barrel cortex layer 2/3, barrel cortex layer 4, dorsolateral striatum, dorsomedial striatum, ventral striatum, basolateral amygdala (BLA), and the CA1 field of the hippocampus. These brain regions were delineated according to the mouse brain atlas of Franklin and Paxinos (2008). Pyramidal neurons were selected from each region, except for barrel cortex layer 4 and the striatum, from which spiny stellate neurons and medium spiny neurons were selected, respectively. Neurons were selected if they were unobscured (by vasculature, glial cells, or other neurons) and appeared to be wholly stained and intact within the section. Entire neurons (excluding axons) were traced at 400× magnification, using a microscope (Zeiss Axioskop 2) with a camera lucida and a 40×/0.75 objective lens. Eight neurons (four per hemisphere) were traced from each region, with the exception of the striatal regions. In these cases, fewer neurons (a minimum of 4, but generally 6-8) were traced from some brains, due to the difficulty in finding neurons in the striatum that were suitable for tracing (i.e., not obscured by vasculature). The mean number of neurons traced per brain from each striatal region was 7.3 for the dorsolateral region, 7.3 for the dorsomedial region, and 6.8 for the ventral region.

Dendritic morphology was assessed by Sholl analysis. For each neuron, a series of concentric circles (with each circle separated by 5 mm in radius) was overlaid over the tracing, centered on the cell soma. The number of intersections between the dendrites and the circles was counted. This count was then converted into real units of length by multiplying by the distance between circles and dividing by the magnification of the tracing. For pyramidal neurons, separate measures of length were calculated for the apical dendritic field (including the primary apical dendrite and any of its oblique dendrites) and the basal dendritic field. The one exception to this was for the BLA pyramidal neurons, in which case we did not differentiate between the apical and basal fields, due to the difficulty in discerning the apical dendrite in many of these cells.

For analysis of spine density, unobscured terminal branches were selected from eight neurons (four per hemisphere), and a segment 10 µm in length or greater, which included the branch tip, was traced at 1000× magnification, using a 100×/1.30 objective. The exception again was the striatal regions, from which branch tips were traced from six neurons (three per hemisphere). For cortical layer 4 spiny stellate neurons and for pyramidal neurons in the BLA, a single terminal branch tip was traced from each neuron. For striatal medium spiny neurons, two terminal branch tips were traced per neuron, with the branches selected from opposite sides of the cell. For all other pyramidal neurons, two branch tips were traced: one from the apical dendritic field and one from the basal field. The number of spines was then counted and divided by the length of the segment to give a measure of spine density. The length of the segment was determined by converting the tracings to digital images and using the “freehand line” tool in ImageJ (http://rsb.info.nih.gov/ij/). The mean ± standard deviation (SD) and median lengths of the dendritic segments were as follows: barrel cortex layer 3, basal: 3.1 ± 0.7, med: 3.0; barrel cortex layer 3, apical: 2.9 ± 0.8, med: 2.9; barrel cortex layer 4: 3.2 ± 0.9, med: 3.1; dorsolateral striatum: 2.3 ± 0.3, med: 2.2; dorsomedial striatum: 2.3 ± 0.3, med: 2.3; ventral striatum: 2.2 ± 0.3, med: 2.2; BLA: 7.8 ± 2.3, med: 7.4; hippocampus, basal: 5.3 ± 1.3, med: 5.3; hippocampus, apical: 5.8 ± 1.5, med: 5.9. No explicit attempt was made to quantify spine density at a fixed distance from the soma, though, in practice, the selection of terminal branches meant that the branch segments analyzed were distal to the soma, at or near the perimeter of the dendritic field.

### 1.2.5 Statistical Analysis

Measures of dendritic length and spine density were averaged across the eight neurons selected from each brain region. Significance testing was conducted by two-way analysis of variance, using IBM SPSS Statistics (Version 24). Statistics are reported in-text below for effects with a *p*-value • .1. All statistics are reported in Supplementary Table S1 (dendritic length) and Table S2 (spine density). Descriptive statistics are reported as mean ± SD.

## 1.3 RESULTS

### 1.3.1 Barrel Cortex

#### 1.3.1.1 Layer 2/3

For the layer 2/3 pyramidal neurons of the barrel cortex (Figure 1A, 1B), the total length of the basal dendritic field (Figure 1C) was significantly larger in ZnT3 KO mice than in WT mice [*F*(1, 23) = 4.51, *p* = .045]. This difference appeared to be driven primarily by the enriched housing groups, though the interaction between genotype and housing was not significant [*F*(1, 23) = 3.03, *p* = .095]. Increased total length of the basal dendritic field could be due to a greater number of branches, longer individual branches, or a combination of both factors. To assess this, the total number of basal dendritic branches was counted. There was no significant effect of genotype, enriched housing, or interaction between genotype and housing, though the ZnT3 KO mice exposed to enriched housing did tend to have more branches than the other three groups (WT-control: 66.9 ± 8.8; KO-control: 66.8 ± 8.5; WT-enriched: 67.3 ± 10.2; KO-enriched: 78.5 ± 9.4). These results suggest that the increased total basal dendritic length in ZnT3 KO mice was most likely due to a combination of more and longer branches. For the apical dendritic field (Figure 1D), the trend was toward greater total length in the ZnT3 KO mice, but the difference between genotypes was not significant [*F*(1, 23) = 3.52, *p* = .073]. Enriched housing significantly decreased the total length of the apical field [*F*(1, 23) = 24.71, *p* < .001], but had no significant effect on the basal dendrites. There was no significant interaction between genotype and housing condition for the apical dendritic field.

**Figure 1.**
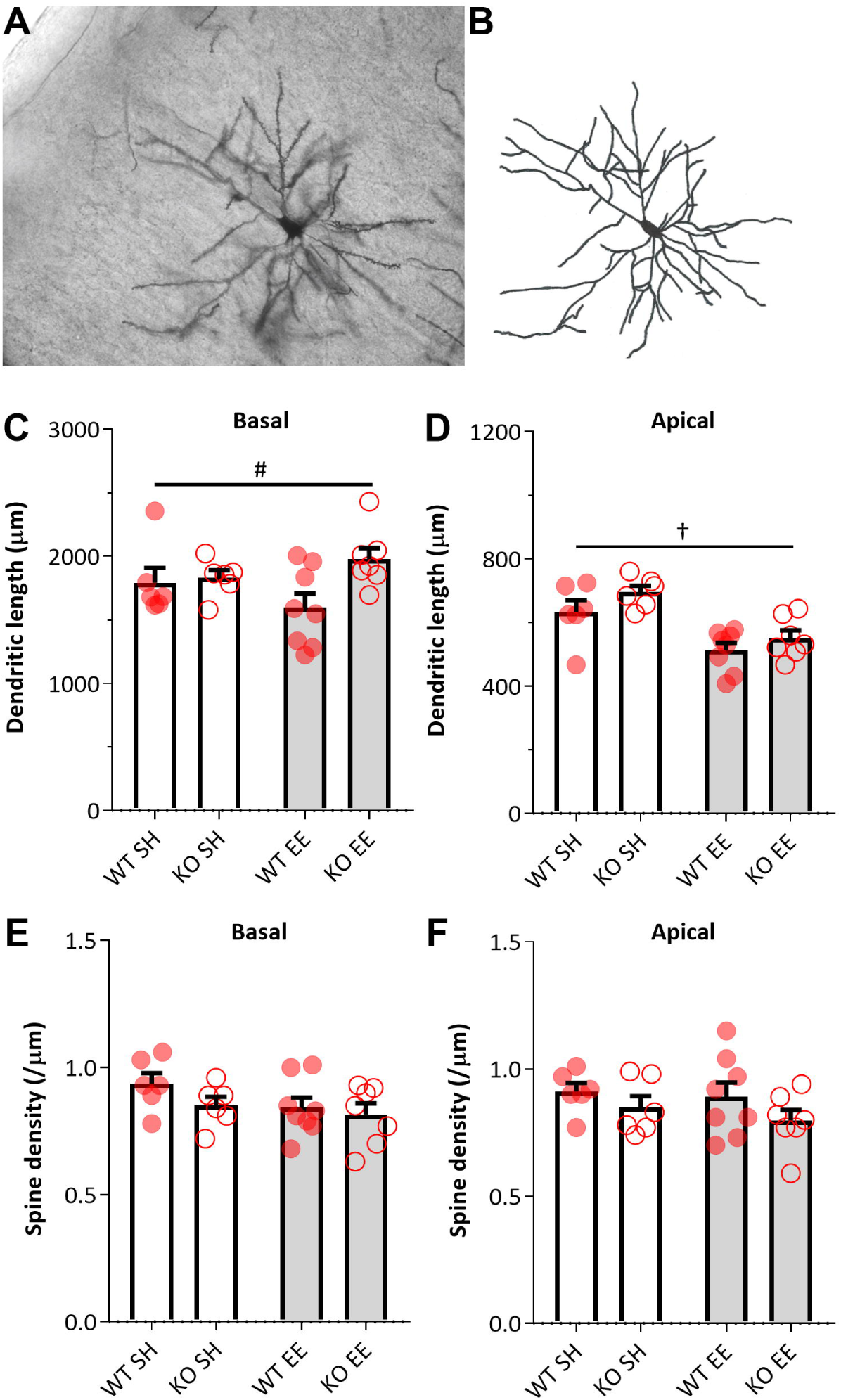
Effects of environmental enrichment and ZnT3 genetic status on the dendritic morphology of layer 2/3 pyramidal neurons in mouse barrel cortex. **(A, B)** Example photomicrograph, and corresponding camera lucida tracing, of a layer 2/3 pyramidal neuron visualized using the Golgi-Cox method. **(C)** The total length of the basal dendritic field was greater in ZnT3 KO mice than in WT mice. Environmental enrichment had no effect on basal dendritic length. **(D)** Environmental enrichment decreased the total length of the apical dendritic field, whereas ZnT3 genotype had no effect on apical dendritic length. **(E, F)** Neither environmental enrichment nor ZnT3 genotype affected the density of spines on the apical or basal dendrites of the layer 2/3 pyramidal neurons in barrel cortex. Spine density was measured on terminal branch tips. Error bars represent 1 standard error of the mean (SEM). SH: standard housing; EE: enriched environment. ^#^ main effect of genotype, *p* < .05; ^†^ main effect of housing condition, *p* < .05

The density of dendritic spines on these neurons did not differ between genotypes for the basal dendrites (Figure 1E). For the apical dendrites (Figure 1F), WT mice tended to have greater spine density than ZnT3 KO mice, though this difference was not significant [*F*(1, 23) = 3.00, *p* = .097]. There was also no effect of enrichment or interaction between genotype and enrichment on apical or basal spine density.

#### 1.3.1.2 Layer 4

For the layer 4 spiny stellate neurons, there was no significant difference in dendritic length (Table 1) or spine density (Table 2) between genotypes, nor was there an effect of environmental enrichment or an interaction between genotype and enrichment on either measure.

**Table 1.** Analysis of dendritic length. Effects of environmental enrichment and ZnT3 genetic status on the dendritic length of neurons across several brain regions in female mice.

| | Dendritic length ($\mu\text{m}$) | | | |
| --- | --- | --- | --- | --- |
|  | Standard housed |  | Enriched environment |  |
| | Wild type<br>($n = 6$) | ZnT3 KO<br>($n = 6$) | Wild type<br>($n = 8$) | ZnT3 KO<br>($n = 7$) |
| **Barrel Cortex** |  |  |  |  |
| Layer 2/3 (basal)<sup>#</sup> | 1793 ± 284 | 1830 ± 146 | 1598 ± 308 | 1979 ± 230 |
| Layer 2/3 (apical)<sup>†</sup> | 634 ± 93 | 695 ± 49 | 514 ± 64 | 551 ± 64 |
| Layer 4 | 903 ± 130 | 951 ± 70 | 826 ± 149 | 905 ± 93 |
| **Striatum** |  |  |  |  |
| Dorsolateral | 1299 ± 150 | 1315 ± 149 | 1243 ± 170 | 1305 ± 188 |
| Dorsomedial | 1000 ± 110 | 1079 ± 140 | 1011 ± 136 | 999 ± 95 |
| Ventral | 783 ± 114 | 704 ± 120 | 819 ± 81 | 815 ± 106 |
| **Amygdala** |  |  |  |  |
| Basolateral nucleus | 1256 ± 344 | 1443 ± 431 | 1075 ± 91 | 1290 ± 225 |
| **Hippocampus** |  |  |  |  |
| CA1 (basal) | 698 ± 106 | 846 ± 159 | 745 ± 104 | 769 ± 134 |
| CA1 (apical) | 609 ± 156 | 764 ± 229 | 693 ± 220 | 738 ± 101 |

Values are reported as mean ± SD.
<sup>#</sup>Effect of genotype, $p < .05$
<sup>†</sup>Effect of housing, $p < .05$

**Table 2.** Analysis of spine density. Effects of environmental enrichment and ZnT3 genetic status on the spine density of neurons across several brain regions in female mice.

| | Spine density (spines/$\mu\text{m}$) | | | |
| --- | --- | --- | --- | --- |
|  | Standard housed |  | Enriched environment |  |
| | Wild type<br>($n = 6$) | ZnT3 KO<br>($n = 6$) | Wild type<br>($n = 8$) | ZnT3 KO<br>($n = 7$) |
| **Barrel Cortex** |  |  |  |  |
| Layer 2/3 (basal) | $0.94 \pm 0.10$ | $0.85 \pm 0.08$ | $0.84 \pm 0.11$ | $0.82 \pm 0.12$ |
| Layer 2/3 (apical) | $0.91 \pm 0.08$ | $0.85 \pm 0.11$ | $0.89 \pm 0.16$ | $0.80 \pm 0.11$ |
| Layer 4 | $0.69 \pm 0.09$ | $0.69 \pm 0.04$ | $0.65 \pm 0.07$ | $0.72 \pm 0.09$ |
| **Striatum** |  |  |  |  |
| Dorsolateral<sup>†</sup> | $0.44 \pm 0.06$ | $0.45 \pm 0.10$ | $0.52 \pm 0.08$ | $0.58 \pm 0.11$ |
| Dorsomedial<sup>†</sup> | $0.37 \pm 0.04$ | $0.44 \pm 0.04$ | $0.47 \pm 0.06$ | $0.48 \pm 0.10$ |
| Ventral<sup>†</sup> | $0.36 \pm 0.10$ | $0.31 \pm 0.06$ | $0.48 \pm 0.10$ | $0.55 \pm 0.05$ |
| **Amygdala** |  |  |  |  |
| Basolateral nucleus | $0.42 \pm 0.09$ | $0.43 \pm 0.07$ | $0.48 \pm 0.11$ | $0.48 \pm 0.07$ |
| **Hippocampus** |  |  |  |  |
| CA1 (basal) | $0.58 \pm 0.08$ | $0.58 \pm 0.08$ | $0.53 \pm 0.11^*$ | $0.55 \pm 0.12$ |
| CA1 (apical) | $0.50 \pm 0.07$ | $0.58 \pm 0.05$ | $0.52 \pm 0.09^*$ | $0.55 \pm 0.11$ |

Values are reported as mean ± SD.
\*For CA1 spine density measures: $n = 7$
<sup>#</sup>Effect of genotype, $p < .05$
<sup>†</sup>Effect of housing, $p < .05$

### 1.3.2 Striatum

#### 1.3.2.1 Ventral

For medium spiny neurons in the ventral striatum (Figure 2A, 2B), there was no effect of genotype on dendritic length, nor was there an interaction between genotype and housing. Enriched housing did tend to increase dendritic length (Figure 2C), though this difference was not significant [*F*(1, 23) = 3.39, *p* = .079]. There was, however, a significant effect of environmental enrichment on spine density in the ventral striatum (Figure 2D), with enrichment increasing the density of spines [*F*(1, 23) = 31.82, *p* < .001]. There was no effect of genotype, or interaction between genotype and housing [*F*(1, 23) = 3.69, *p* = .067], on spine density, though this interaction was close to the cut-off for significance, with enrichment tending to increase spine density to a greater degree in ZnT3 KO mice than in WT mice.

**Figure 2.**
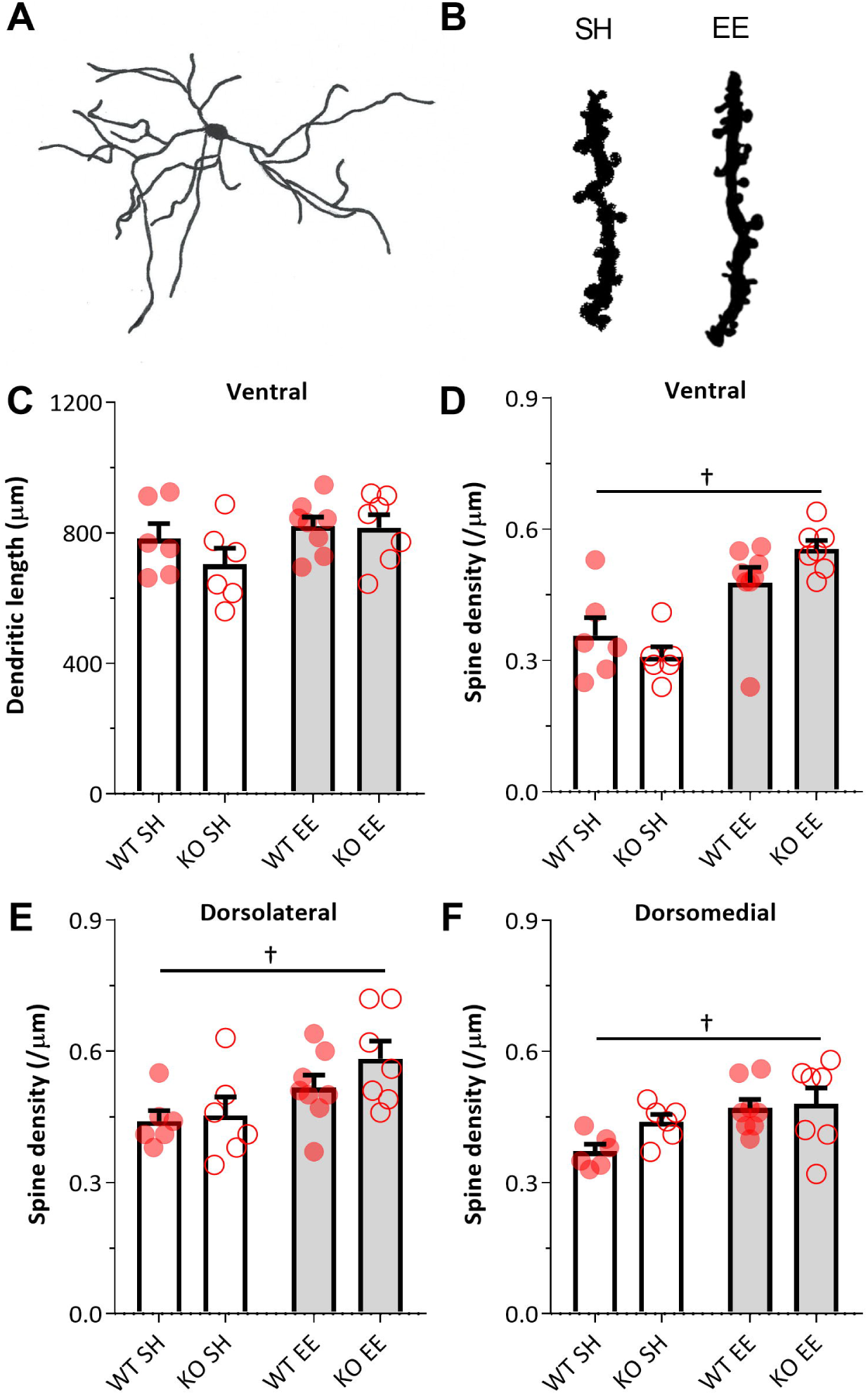
Effects of environmental enrichment and ZnT3 genetic status on the dendritic morphology of medium spiny neurons in the mouse striatum. **(A)** Example camera lucida tracing of a medium spiny neuron in the ventral striatum. **(B)** High magnification tracings, used to quantify spine density, of terminal dendritic segments from ventral striatum medium spiny neurons, from a mouse in standard housing (SH) and a mouse exposed to environmental enrichment (EE). **(C)** Neither environmental enrichment nor ZnT3 genotype affected the dendritic length of medium spiny neurons in any of the three striatal compartments examined, including the ventral striatum, for which data are displayed here. **(D, E, F)** Environmental enrichment did, however, increase the density of spines on the terminal branch tips of medium spiny neurons in the ventral, dorsolateral, and dorsomedial regions of the striatum. Error bars represent 1 SEM. ^†^ main effect of housing condition, *p* < .05

#### 1.3.2.2 Dorsolateral and Dorsomedial

In the dorsolateral and dorsomedial striatum, there was no effect of genotype or enriched housing on dendritic length (Table 1), nor was there an interaction between genotype and enriched housing. As in the ventral striatum, enriched housing increased spine density on striatal neurons in both the dorsolateral region [*F*(1, 23) = 9.00, *p* = .006; Figure 2E] and the dorsomedial region [*F*(1, 23) = 8.15, *p* = .009; Figure 2F]. There was no effect of genotype, or interaction between genotype and housing condition, on spine density in either region.

### 1.3.3 Amygdala

For the pyramidal neurons in the basolateral nucleus of the amygdala, there was no significant effect of enriched housing on dendritic length, nor was there an interaction between genotype and housing (Table 1). Dendritic length tended to be greater in ZnT3 KO mice compared to WT mice, though this difference was not significant [*F*(1, 23) = 3.27, *p* = .084]. Similarly, spine density (Table 2) was not affected by genotype or enriched housing, and there was no interaction between genotype and enriched housing.

### 1.3.4 Hippocampus

Finally, dendritic length (Table 1) and spine density (Table 2) were assessed for the pyramidal neurons in the CA1 field of the hippocampus. For the total length of the basal dendritic field, there was no effect of genotype or enriched housing, nor was there an interaction between genotype and housing. Basal dendritic length did tend to be greater in the ZnT3 KO mice than in the WT mice [*F*(1, 23) = 3.14, *p* = .090], although this difference was not significant. For apical dendritic length, there was no effect of genotype or enriched housing, nor was there an interaction between these two factors.

For the density of spines on both the basal dendrites and apical dendrites of the CA1 pyramidal neurons, there was no effect of genotype or enriched housing, nor was there an interaction between genotype and housing. Hippocampal spine density could not be assessed for one brain (WT-EE) due to a broken slide.

## 1.4 DISCUSSION

The present study was undertaken to address two questions. The first was whether elimination – during development and throughout life – of ZnT3, the vesicular zinc transporter, would impact neuronal morphology in several brain regions that are normally richly innervated by zinc-containing axon terminals. We found that it generally does not; neuronal morphology in female ZnT3 KO mice (males were not assessed in the present experiment) was normal in the striatum, amygdala, and hippocampus. The one exception was pyramidal neurons in the barrel cortex, in which the total length of the basal dendritic arbour was greater in ZnT3 KO mice compared to WT animals. Our results indicate that the greater dendritic length was most likely due to a combination of these neurons having both more and longer dendritic branches in ZnT3 KO mice compared to WT mice, as an increase in dendritic branch number alone was not sufficient to account for the significant difference between genotypes.

The second question was whether ZnT3 KO mice would exhibit alterations in neuronal morphology in response to environmental enrichment. Given evidence that vesicular zinc may be involved in experience-dependent plasticity in the neocortex (Czupryn & Skangiel-Kramska, 2001; Brown & Dyck, 2002, 2005; Nakashima & Dyck, 2010; Kalappa et al., 2015) and hippocampus (Suh et al., 2009), we hypothesized that mice lacking vesicular zinc would show fewer or smaller effects of enrichment relative to WT mice. Our results provided little support for this hypothesis. While enrichment affected aspects of neuronal morphology in the barrel cortex and striatum, none of these effects was exclusive to, or significantly larger in, WT mice compared to ZnT3 KO mice. Indeed, the closest case to an interaction between genotype and environmental enrichment was in spine density on neurons in the ventral striatum, and it was ZnT3 KO mice that tended to show the larger effect, though not significantly so. Thus, it appears that vesicular zinc, while important for some forms of experience-dependent plasticity, is not required for experience-dependent alterations in neuronal morphology induced by environmental enrichment – at least under the conditions and in the regions examined in the present study.

The general lack of morphological abnormalities identified in ZnT3 KO mice is not entirely surprising, considering that these mice exhibit no pronounced behavioural phenotype (Cole et al., 2001; but see Yoo et al., 2016) but instead show mild abnormalities or deficiencies across several behavioural domains (Martel et al., 2010, 2011; Sindreu et al., 2011; Thackray et al., 2017; Wu & Dyck, 2018, Kumar et al., 2019). This suggests that most behavioural abnormalities in these mice are mediated not by gross changes in neuronal morphology, but by changes on a smaller scale at the level of neurotransmission, which vesicular zinc has been shown to modulate through its effects on several different receptors (Perez-Rosello et al., 2013; Anderson et al., 2015; Kalappa et al., 2015). The exception, based on our present findings, might be the impairment in whisker-mediated texture discrimination (Wu & Dyck, 2018), which could be related to the morphological abnormality we observed in the barrel cortex, given that texture discrimination is a barrel-cortex dependent function (Guic-Robles et al., 1992). To speculate, it is possible that an aberrant increase in dendritic length could result in a reduction in the specificity of the sensory input received by pyramidal neurons in barrel cortex, contributing to an impairment in discriminating between fine differences in texture. In addition to following up on this possibility, it would be interesting to examine whether our observation of abnormal morphology in barrel cortex neurons extends to other sensory areas such as auditory cortex, especially given that ZnT3 KO mice also exhibit deficits in auditory frequency discrimination (Kumar et al., 2019).

Our observation of dendritic hypertrophy in the barrel cortex is noteworthy for an additional reason. Previously, we conducted an MRI volumetric analysis of several brain regions in male ZnT3 KO mice and found that the volume of parietal cortex (or, more precisely, a subsection of parietal cortex) was increased by 3.6% (McAllister et al., 2018). Specifically, cortical volume was measured around the position along the anterior-posterior axis where the dorsal hippocampus is situated, which includes the area from which barrel cortex neurons were traced in the present study. This effect is also supported by an MRI analysis conducted by Yoo et al. (2016), who examined volume over a broader area of neocortex, and found it to be larger in male ZnT3 KO mice by ∼5% compared to WT mice. Increased cortical volume could potentially reflect many different aspects of the underlying anatomy (e.g., changes in neurons, glial cells, or vasculature), which cannot be differentiated by MRI. Our present analysis suggests that an increase in the length or number of dendritic processes emanating from cortical pyramidal neurons may be the cause – or at least one contributing factor. Further results by Yoo et al. (2016) suggest other factors. They found – again, in males – that ZnT3 KO mice show increased cortical expression of NeuN, a marker for neurons, as well as an increase of ∼12% in the number of NeuN-positive cells in the region of dorsal cortex appearing to correspond to the restrosplenial and parietal association areas (see their figure 2d). Their results also show increased expression of neurofilament H, which is suggestive of an increase in the length or number of neuronal processes, particularly axons. It should be noted though, that these findings were observed at 5 weeks of age, so it is uncertain if they would persist into maturity.

The results of Yoo et al. (2016) also provide a potential mechanism underlying hypertrophic brain development in ZnT3 KO mice: increased levels of brain-derived neurotrophic factor (BDNF) in the neocortex and hippocampus – an observation which is consistent with previous findings of increased hippocampal BDNF levels in adult ZnT3 KO mice (Helgager et al., 2014). The effects of BDNF on the dendritic morphology of neocortical and hippocampal neurons are complex and dependent on a number of factors (McAllister et al., 1997; Horch et al., 1999; Murphy et al., 1998; Horch & Katz, 2002), but there are cases where BDNF acts to promote dendritic growth. Particularly relevant is that the total dendritic length and complexity of dentate granule cells is increased in mice that chronically overexpress BDNF (Tolwani et al., 2002). Thus, it is possible that an excess of BDNF in ZnT3 KO mice is responsible for the increased dendritic branching of layer 2/3 pyramidal neurons in the somatosensory cortex – though it is unclear why this effect would not also apply to pyramidal cells in the hippocampus. It should be noted, however, that we and others have failed to detect increased BDNF levels in young or mature ZnT3 KO mice (Adlard et al., 2010; McAllister et al., submitted), so this potential mechanism requires further verification and investigation.

In addition to the effects of genetically eliminating vesicular zinc, we observed that 8 weeks of environmental enrichment significantly altered the morphology of neurons in both the barrel cortex and striatum. Regarding the former, exposure to enriched housing decreased the total length of the apical dendritic arbour in layer 2/3 pyramidal neurons. This was somewhat surprising, considering that environmental enrichment has a generally expansionary effect on dendritic length in cortical neurons (Greenough & Volkmar, 1973; Greenough et al., 1973; Uylings et al., 1978; Kolb et al., 2003a; Kolb et al., 2003b; Gelfo et al., 2009). Interestingly, previous research on rats has shown that enriched housing in a naturalistic habitat causes shrinking, sharpening, and weakening of the whisker receptive fields in the supragranular layers of the barrel cortex (Polley et al., 2004). It is possible, then, that the reduction in dendritic length observed in the present experiment may be related to this functional refinement in the vibrissal somatotopic map induced by enrichment, though this potential link is merely correlational and would require further investigation to substantiate.

The most robust morphological effect of enrichment that we observed was on the medium spiny neurons of the striatum. Specifically, environmental enrichment increased the density of spines on these cells in all three striatal regions – dorsolateral, dorsomedial, and ventral – that were examined. This is consistent with previous findings by Comery et al. (1995, 1996) and Kolb et al. (2003a) of increased spine density on striatal medium spiny neurons following environmental enrichment in rats. In addition, Kolb et al. (2003a) reported an increase in the overall dendritic length of medium spiny neurons in the ventral striatum, which we did not observe in mice. A previous experiment also using mice was similarly unable to detect an effect of environmental enrichment on dendritic length of spiny neurons in the ventrolateral striatum (Faherty et al., 2003), so it is possible that the discrepancy with the results of Kolb et al. (2003a) was due to a species difference. Alternatively, the cause could be differences in the age at which animals were exposed to environmental enrichment or the length of time spent in the enriched environment; Kolb et al. (2003a) of increased spine density on striatal medium spiny neurons following environmental enrichment in rats. In addition, Kolb et al. (2003a) used a longer period of enrichment and also initiated the enrichment in adulthood, whereas our enrichment procedure began at 4 weeks of age. We opted to initiate enrichment shortly after weaning, as this is fairly common practice in research on the morphological effects of environmental enrichment, including in many of the classic studies on the topic (e.g., Diamond et al., 1972, Greenough et al., 1973; Volkmar & Greenough, 1973). It is possible that the effects of enrichment may interact with developmental processes or hormonal changes relating to the onset of puberty, which could influence the outcomes of environmental enrichment in studies, such as the present one, where enrichment is initiated at or shortly after weaning. Indeed, Kolb et al. (2003b) compared the effects of 3 months of environmental enrichment beginning at either weaning or adulthood in rats, and found that some of the effects of enrichment on adult rats were eliminated – or even reversed – in rats that were introduced to an enriched environment as weanlings. Thus, we cannot rule out that we may have observed different results if we initiated environmental enrichment in adulthood in our experiment.

In general, it is noteworthy that we observed few effects of environmental enrichment, relative to what has been described by previous researchers studying the effects of enrichment on rats (e.g., Kolb et al., 2003a, 2003b; Gelfo et al., 2009). In addition to the methodological factors already highlighted (i.e., species differences, age at onset of enrichment), there are at least two other explanations. First, previous work has shown that when rats are housed in enriched environments, the largest effects – at least in terms of cortical depth and weight – are seen after 30 days (Rosenzweig et al., 1969; Diamond et al., 1972). By 80 days, the effects are somewhat diminished. In the present experiment, we used an 8-week (56-day) period of enrichment, based on previous studies that observed effects on neuronal morphology in mice after 40-60 days of enriched housing (Rampon et al., 2000; Restivo et al., 2005; Karelina et al., 2012). It is possible that we would have observed more or larger effects if we investigated neuronal morphology after a shorter period of enrichment. Second, there is some evidence that female rats show fewer effects of environmental enrichment than males, at least in cortical neurons (Juraska, 1984; Kolb et al., 2003b) – although the opposite seems to be the case in the hippocampus (Juraska et al., 1985, 1989) – so it is possible that the use of female mice in the present experiment resulted in fewer effects of enrichment on neuronal morphology. Again, it could be the case that a species difference is the cause of the discrepancy between the current finding that enrichment had no effect on hippocampal neurons and the previous results from Juraska et al. (1985, 1989) that enrichment increased dendritic length in hippocampal neurons. Alternatively, the discrepancy could be due to differences in the hippocampal sub-regions from which neurons were sampled (CA1 in our study vs. CA3 or dentate gyrus) or differences in the control condition that was used (standard housing in our study vs. isolated housing).

On the topic of sex differences, a limitation of the present experiment was the use of only female mice. To provide social enrichment, our enrichment protocol involved the group-housing of mice from multiple, similarly-aged litters. Compared to female mice, males are much more likely to fight and injure one another, especially when they are group-housed with non-littermates (Deacon, 2011). Beyond the obvious concerns about animal welfare, this also raised concerns about the potentially confounding effects of stress, particularly because stress is well-established to alter neuronal morphology in many of the same brain regions examined in the present study, such as the amygdala, hippocampus, and ventral striatum (Watanabe et al., 1992; Magariños et al., 1996; Sousa et al., 2000; Vyas et al., 2004; Bessa et al., 2013). For this reason, we chose to use only female mice in this experiment. The obvious drawback is that we cannot conclude whether our results would extend to males – though given the apparent consistency, discussed above, between our present results in females and our MRI results from males, it seems quite possible that at least some of them would.

In summary, the present study was conducted to determine whether mice that lack ZnT3 and vesicular zinc exhibit abnormalities in neuronal morphology, with the additional objective of examining whether these mice exhibit experience-dependent changes in neuronal morphology in response to housing in an enriched environment. We found that the dendritic morphology of neurons in these mice was broadly normal across several brain regions, including the primary somatosensory cortex (barrel field), striatum, amygdala, and hippocampus. The exception was the pyramidal neurons in layer 2/3 of the barrel cortex, in which basilar dendritic length was greater in ZnT3 KO mice, consistent with recent findings of increased cortical volume in these animals. We also found that elimination of ZnT3 neither augmented nor diminished the effects of environmental enrichment. Independently of genotype, environmental enrichment exerted a few effects on neuronal morphology, some of which were consistent with previous findings in mice and rats (i.e., an increased density of spines on striatal neurons), although, to our knowledge, the finding that enrichment decreased apical dendritic length in layer 2/3 neurons of the barrel cortex is novel. In conclusion, the present results are consistent with the dominant narrative of the past literature on ZnT3 KO mice, which is that a lack of ZnT3 and vesicular zinc throughout life does not produce a pronounced behavioural or neuroanatomical phenotype, though it does cause subtle abnormalities. Furthermore, while ZnT3 KO mice are deficient in some forms of experience-dependent plasticity, the present results provide no evidence that this applies to changes in the morphology of dendrites and dendritic spines in response to environmental enrichment.

## 1.5 ACKNOWLEDGEMENTS

This work was supported by the Natural Sciences and Engineering Research Council of Canada (Discovery Grant and doctoral scholarship); the Killam Trusts (doctoral scholarship); and Mitacs (Globalink Research Awards). The funding sources had no role in the design of this study; in the collection, analysis, and interpretation of data; in the writing of the report; or in the decision to submit this article for publication.

## Footnotes

* BDNF: brain-derived neurotrophic factor; BLA: basolateral amygdala; CNS: central nervous system; KO: knockout; MRI: magnetic resonance imaging; ZnT3: zinc transporter 3; WT: wild type.

