## Supplemental Tables for "Effects of enriched housing on the neuronal morphology of mice that lack zinc transporter 3 (ZnT3) and vesicular zinc"

SUPPLEMENTARY MATERIAL

| **Table S1. ANOVA results for dendritic length.** | | | |
| --- | --- | --- | --- |
|  | Dendritic length | | |
|  | Genotype | Environment | Interaction |
| **Barrel Cortex** |  |  |  |
| Layer 2/3 (basal) | ***F* = 4.51, *p* = .045** | *F* = 0.06, *p* = .815 | *F* = 3.03, *p* = .095 |
| Number of branches | *F* = 2.34, *p* = .140 | *F* = 2.79, *p* = .108 | *F* = 2.38, *p* = .136 |
| Layer 2/3 (apical) | *F* = 3.53, *p* = .073 | ***F* = 24.71, *p* < .001** | *F* = 0.21, *p* = .649 |
| Layer 4 | *F* = 1.96, *p* = .175 | *F* = 1.84, *p* = .189 | *F* = 0.12, *p* = .737 |
| **Striatum** |  |  |  |
| Dorsolateral | *F* = 0.37, *p* = .551 | *F* = 0.27, *p* = .607 | *F* = 0.13, *p* = .725 |
| Dorsomedial | *F* = 0.49, *p* = .491 | *F* = 0.53, *p* = .473 | *F* = 0.94, *p* = .344 |
| Ventral | *F* = 1.05, *p* = .317 | *F* = 3.39, *p* = .079 | *F* = 0.85, *p* = .367 |
| **Amygdala** |  |  |  |
| Basolateral nucleus | *F* = 3.27, *p* = .084 | *F* = 2.28, *p* = .145 | *F* = 0.02, *p* = .899 |
| **Hippocampus** |  |  |  |
| CA1 (basal) | *F* = 3.14, *p* = .090 | *F* = 0.09, *p* = .763 | *F* = 1.67, *p* = .209 |
| CA1 (apical) | *F* = 1.93, *p* = .179 | *F* = 0.16, *p* = .698 | *F* = 0.59, *p* = .451 |
| Degrees of freedom for all tests: 1, 23. | | | |

| **Table S2. ANOVA results for spine density.** | | | |
| --- | --- | --- | --- |
|  | Spine density | | |
|  | Genotype | Environment | Interaction |
| **Barrel Cortex** |  |  |  |
| Layer 2/3 (basal) | *F* = 1.94, *p* = .177 | *F* = 2.51, *p* = .127 | *F* = 0.52, *p* = .479 |
| Layer 2/3 (apical) | *F* = 3.00, *p* = .097 | *F* = 0.59, *p* = .452 | *F* = 0.10, *p* = .759 |
| Layer 4 | *F* = 1.94, *p* = .177 | *F* = 0.03, *p* = .858 | *F* = 1.87, *p* = .185 |
| **Striatum** |  |  |  |
| Dorsolateral | *F* = 1.39, *p* = .250 | ***F* = 9.00, *p* = .006** | *F* = 0.55, *p* = .466 |
| Dorsomedial | *F* = 2.33, *p* = .141 | ***F* = 8.15, *p* = .009** | *F* = 1.30, *p* = .267 |
| Ventral | *F* = 0.19, *p* = .669 | ***F* = 31.82, *p* < .001** | *F* = 3.69, *p* = .067 |
| **Amygdala** |  |  |  |
| Basolateral nucleus | *F* = 0.01, *p* = .921 | *F* = 2.40, *p* = .135 | *F* = 0.11, *p* = .744 |
| **Hippocampus** |  |  |  |
| CA1 (basal) | *F* = 0.07, *p* = .796 | *F* = 1.17, *p* = .292 | *F* = 0.06, *p* = .805 |
| CA1 (apical) | *F* = 2.27, *p* = .146 | *F* = 0.03, *p* = .867 | *F* = 0.44, *p* = .515 |
| Degrees of freedom for all tests: 1, 23; except for CA1 spine density (basal and apical): 1, 22. | | | |
